# Small-molecule targeting MuRF1 enhances functional exercise capacity in rats: an exploratory study

**DOI:** 10.64898/2026.02.05.704003

**Authors:** Fernando Ribeiro, Luis D. Chinait, Maria Rita C. Rodrigues, Siegfried Labeit, Anselmo S. Moriscot

**Affiliations:** Department of Anatomy, Institute of Biomedical Sciences, University of São Paulo, 05508-000 São Paulo, Brazil; Department for Integrative Pathophysiology, DZHK Partner Site Mannheim-Heidelberg, Medical Faculty Mannheim, University of Heidelberg, 68169 Mannheim, Germany; Myomedix GmbH, 69151 Neckargemünd, Germany

**Author notes:** Corresponding Author: Anselmo S. Moriscot, Department of Anatomy, Institute of Biomedical Sciences, University of São Paulo.

**Keywords:** Skeletal muscle, aerobic exercise capacity, muscle histology, ubiquitin-proteasome system, TRIM63

## Abstract

Maintenance of skeletal muscle function is essential for functional independence, quality of life and healthspan. Muscle RING-finger protein-1 (MuRF1) negatively regulates muscle function and mass through ubiquitination and degradation of muscle proteins. Accordingly, genetic and pharmacological inhibition of MuRF1 attenuates muscle wasting and weakness under catabolic stress. To explore the potential of MuRF1 inhibitors (e.g., MyoMed-205) to improve muscle health, we investigated here the long-term effects of MyoMed-205 on functional capacity and muscle physiology in rats under basal conditions. Wistar rats were randomized to control or MyoMed-205 groups and were followed for 4 or 8 weeks. Body weight, food and water intake, and exercise capacity were monitored weekly. At each endpoint, the soleus muscle was collected for histological analyses. MyoMed-205-treated rats showed normal basic survival-related behaviors and body growth. After 8 weeks, MyoMed-205-treated animals exhibited enhanced exercise capacity (speed (m/min): +45%, p = 0.01; endurance (min): +47%, p = 0.03; and distance covered (m): +87%, p = 0.04) compared with baseline performance. Conversely, no differences were found in soleus fiber type distribution, cross-sectional area, or lipid and collagen content. Our findings indicate that MyoMed-205 enhances functional exercise capacity independently of changes in soleus muscle structure in rats under basal conditions.

## Introduction

Skeletal muscles comprise approximately 40% of the human body mass and serve important physiological roles beyond locomotion, including breathing, amino acid storage, systemic metabolism, thermogenesis, hemodynamic and endocrine functions ^1^. Adequate development and maintenance of muscle mass and function are associated with increased functional independence, quality of life and healthspan. Several catabolic conditions (e.g., immobilization, sepsis, cancer, heart failure, neuromuscular dystrophies, mechanical ventilation and ageing) promote muscle wasting and weakness. Furthermore, loss of muscle mass and strength correlates with increased risks of comorbidity and mortality ^2–4^. In this sense, skeletal muscle atrophy and dysfunction result from complex biological processes comprising distinct triggering factors, such as neural, mechanical, metabolic and hormonal mechanisms ^5^.

Muscle RING-finger protein-1 (MuRF1) is encoded by the TRIM63 gene. MuRF1 functionally acts as an ubiquitin E3 ligase known to be involved in the control of skeletal muscle function, mass and metabolism (for review, see ^6,7^). MuRF1 was initially characterized over 20 years ago and has since then been recognized as a key biomarker of muscle wasting in several catabolic states ^8,9^. Interestingly, MuRF1 gene knockout attenuates muscle wasting and weakness under distinct stress models such as denervation, mechanical unloading, or myopathies ^6,10–12^. Under basal conditions, MuRF1 inactivation contributes to a positive protein net balance linked to the activation of anabolic pathways (e.g., Akt-mTOR), and the downregulation of proteolytic systems (e.g., ubiquitin-proteasome) ^13,14^. These findings suggest that MuRF1 may also play an important role in the control of muscle mass and function under basal conditions.

The recent development of pharmacological strategies for targeting MuRF1 has opened new opportunities to counteract muscle wasting and weakness during musculoskeletal disorders, while also enabling deeper investigation of underlying molecular mechanisms. In this context, previous experimental studies demonstrated that MuRF1 inhibition attenuates skeletal muscle atrophy and contractile dysfunction in conditions such as heart failure ^15–18^, cancer cachexia, chemotherapy ^19,20^, as well as acquired myopathies linked to corticosteroids ^15^ and diabetes ^21^. Most recently, our group showed that MyoMed-205, a small-molecule targeting MuRF1, protects against muscle wasting and weakness under acute stress conditions, such as denervation-induced diaphragm dysfunction and atrophy ^22,23^. Collectively, these findings suggest that pharmacological strategies targeting MuRF1, such as MuRF1 inhibitors, hold significant translational potential to protect muscle mass and function under catabolic or disease conditions.

Previous studies show that abnormal regulation of MuRF1 expression could be associated with impaired physical capacity and frailty, while interventions known to enhance muscle health, such as chronic exercise training, reportedly downregulate MuRF1 ^24,25^. However, the effects of MuRF1 inhibitors on muscle physiology and their potential for improving muscle health under basal conditions have not been studied previously. In this exploratory study, we investigated the time-dependent effects of MyoMed-205, a small-molecule targeting MuRF1, on body growth, functional capacity, and skeletal muscle structure in rats. We hypothesize that long-term MyoMed-205 administration would improve overall functional capacity and muscle mass under basal conditions.

## Material and Methods

### Animals and Ethical Considerations

Wistar male rats (∼1-2 months old) were used in this study. The animals were kept in standard cages under controlled environmental conditions (24 ± 1°C, 12 h/12 h light-dark cycle) with access to food and water *ad libitum*. This study was approved and followed the institutional guidelines for animal care and use for research (CEUA ICB-USP #5143091020).

### Study Design

Twelve Wistar rats were initially randomized into two experimental groups: Control (CON, n = 6) and MyoMed-205 (205, n = 6). These animals were monitored over 4 or 8 weeks, respectively, ultimately resulting in four groups (n = 3): Control 4 weeks (CON_4w), MyoMed-205 4 weeks (205_4w), Control 8 weeks (CON_8w), and MyoMed-205 8 weeks (205_8w), respectively. Rat body weight, food and water intake, as well as exercise capacity, were monitored weekly. After each endpoint (i.e., at the 4^th^ and 8^th^ weeks), the rats were euthanized by exsanguination under anesthesia and analgesia induced by ketamine and xylazine mix. Immediately after, the soleus muscle was collected, quickly snap-frozen in 2-methylbutane chilled in liquid nitrogen, and stored at −80°C until further analysis.

### Standard and MyoMed-205 Enriched Rodent Food

To investigate the long-term effects of the small-molecule targeting MuRF1, the MyoMed-205 compound was incorporated with standard rodent food chow (MyoMed-205 0.2% w/w) for oral delivery at a saturating level, similar to previous investigations ^19,21^, for this long-term study (Rhoster #RH29586 modified, Brazil). Control animals had access to the unmodified standard rodent food chow (Rhoster #RH29586, Brazil).

### Exercise Capacity Test

Exercise capacity was determined using an incremental treadmill ramp protocol. After a one-week familiarization period for all animals, the test was performed using a motorized rodent treadmill (AVS Projects, São Carlos, Brazil) set at 0° inclination. The protocol started with an initial speed of 5 m/min, and incremental increases of 5 m/min every 3 minutes were made until the animals reached physical exhaustion. Throughout the test, rats were encouraged to maintain running pace through gentle manual stimulation at the rear of the treadmill lane when appropriate. Physical exhaustion was defined as the inability of the animal to maintain a running state despite receiving at least three consecutive external stimuli within 30 seconds. Immediately upon reaching physical exhaustion, peak running speed (m/min), exercise duration (min), and total distance covered (m) were recorded as indices of maximal exercise capacity. Exercise capacity tests were conducted weekly by the same operator and during the same time of day (5:00 – 6:00 pm) for all animals and groups to minimize inter-subject and circadian-related variability in exercise capacity measurements.

### Hematoxylin and Eosin Staining

For qualitative histological analysis, soleus muscle cryosections (10 µm thick) were prepared using a cryostat at −20°C (Leica CM1850, Germany) and subsequently staining with hematoxylin and eosin (H&E). For this, muscle cryosections were allowed to thermoequilibrate for 30 minutes at room temperature. Next, samples were stained with Hematoxylin (Amresco, #0701, USA) for 10 minutes and rinsed three times with distilled water. Next, samples were stained with Eosin (Amresco, #0109, USA) for 2 minutes and rinsed three times with distilled water. Next, samples were incubated in a series of graded ethanol solutions (70%, 95% and 100%) for 2 minutes each. After this, samples were incubated in a two-series of xylene solutions for 2 minutes each. The H&E preparations were finished using a mounting medium (Permount, Fisher Chemical #SP15) and a coverslip. Photomicrographs were acquired using the Axio Scope A1 microscope (Zeiss, Germany).

### Immunohistochemistry for Muscle Fiber Typing and Morphometry

Soleus muscle cryosections (10 µm thick) were allowed to thermoequilibrate at room temperature for 30 minutes. Samples were washed with PBS solution (1 x 5 minutes) and incubated with M.O.M. blocking reagent (MKB-2213-1, VectorLabs) at room temperature for 1 hour. Next, samples were washed with PBS (3 x 5 minutes) and incubated overnight with primary antibodies at 4°C (anti-MHC I (#BA-F8 DSHB, mouse IgG2b, 1:300), anti-MHC IIa (#SC-71 DSHB, mouse IgG1, 1:300), and anti-dystrophin (SC#15376, rabbit IgG, 1:500). In the following day, tissues were washed with PBS (3 x 5 minutes) and incubated with secondary antibodies at 37°C for 1 hour (Alexa Fluor 647 goat anti-mouse IgG2b (Jackson ImmunoResearch, West Grove, PA, USA, #115-607-187, 1:300), Cy2 goat anti-mouse IgG1 (Jackson ImmunoResearch #115-225-205, 1:300), and Cy3 goat anti-rabbit IgG (Jackson ImmunoResearch #111.165.003, 1:200). Next, samples were washed with PBS (3 x 5 minutes) and the histological slides were finished with mounting medium and a coverslip. Photomicrographs were acquired using the Axio Scope A1 microscope (Zeiss, Germany). Muscle fiber type distribution and cross-sectional area (CSA) analysis were performed using ImageJ (Fiji, NIH, USA). Fiber type distribution was determined by the relative distribution of fiber types obtained from whole muscle fiber analysis, while fiber CSA was determined as the mean of 300 fiber CSA measurements per sample.

### Oil Red Staining

Soleus muscle cryosections (10 µm thick) were allowed to thermoequilibrate at room temperature for 30 minutes. Next, samples were fixed using 10% PFA solution for 10 minutes and rinsed with distilled water and 60% isopropyl alcohol solutions. Next, samples were incubated with Oil Red O (ORO) (Sigma #O0635) working solution for 30 minutes and rinsed with distilled water and 60% isopropyl alcohol solutions. ORO-stained histological preparations were finished using glycerine jelly aqueous-based mounting medium and a coverslip. Photomicrographs were acquired using the Axio Scope A1 microscope (Zeiss, Germany). Representative mean of soleus muscle intracellular lipid content was determined by the assessment of 100 muscle fibers per sample using 400x magnification photomicrographs.

### Picrosirius Staining

Soleus muscle cryosections (10 µm thick) were allowed to thermoequilibrate at room temperature for 30 minutes. Next, samples were fixed using 4% PFA solution for 15 minutes and rinsed three times with distilled water. Next, samples were incubated in Picrosirius red solution (0.1% in picric acid) for 30 minutes and rinsed with 0.5% acetic acid in distilled water solution. Next, samples were rinsed in a graded ethanol solution series (70%, 95%, and 100%) for 30 seconds each. Next, samples were incubated with a two-series of xylene solutions for 2 minutes each. Finally, the histological preparations were finished using a mounting medium (Permount, Fisher Chemical #SP15) and a coverslip. Photomicrographs were acquired using the Axio Scope A1 microscope (Zeiss, Germany). Muscle collagen content was determined by the quantification of the relative collagen area out of the whole muscle sample area for each sample using 50x magnification photomicrographs.

### Statistical Analysis

Data processing and analysis were performed with R (Version 2024.09.0+375). The Shapiro-Wilk test was used to assess data normality. Welch’s t-test was used for comparisons between two groups for parametric data without assuming equal variances. Alternatively, the non-parametric Mann-Whitney test was considered for comparisons between two groups if appropriate. Data are presented as the mean ± standard deviation. Statistical significance was accepted as p < 0.05.

## Results

### MyoMed-205 does not affect basic survival-related behaviors, body and vital organs weight after 8 weeks of administration in rats

To assess whether long-term MyoMed-205 administration affects rat development, we weekly monitored body weight, food and water intake over 8 weeks (Figure 1A). MyoMed-205-treated rats exhibit normal body weight development over time compared with controls (Figure 1B). Consistently, basic survival-related behaviors such as food and water intake dif not differ between MyoMed-205-treated and control animals (Figure 1C-1D).

**Figure 1.**
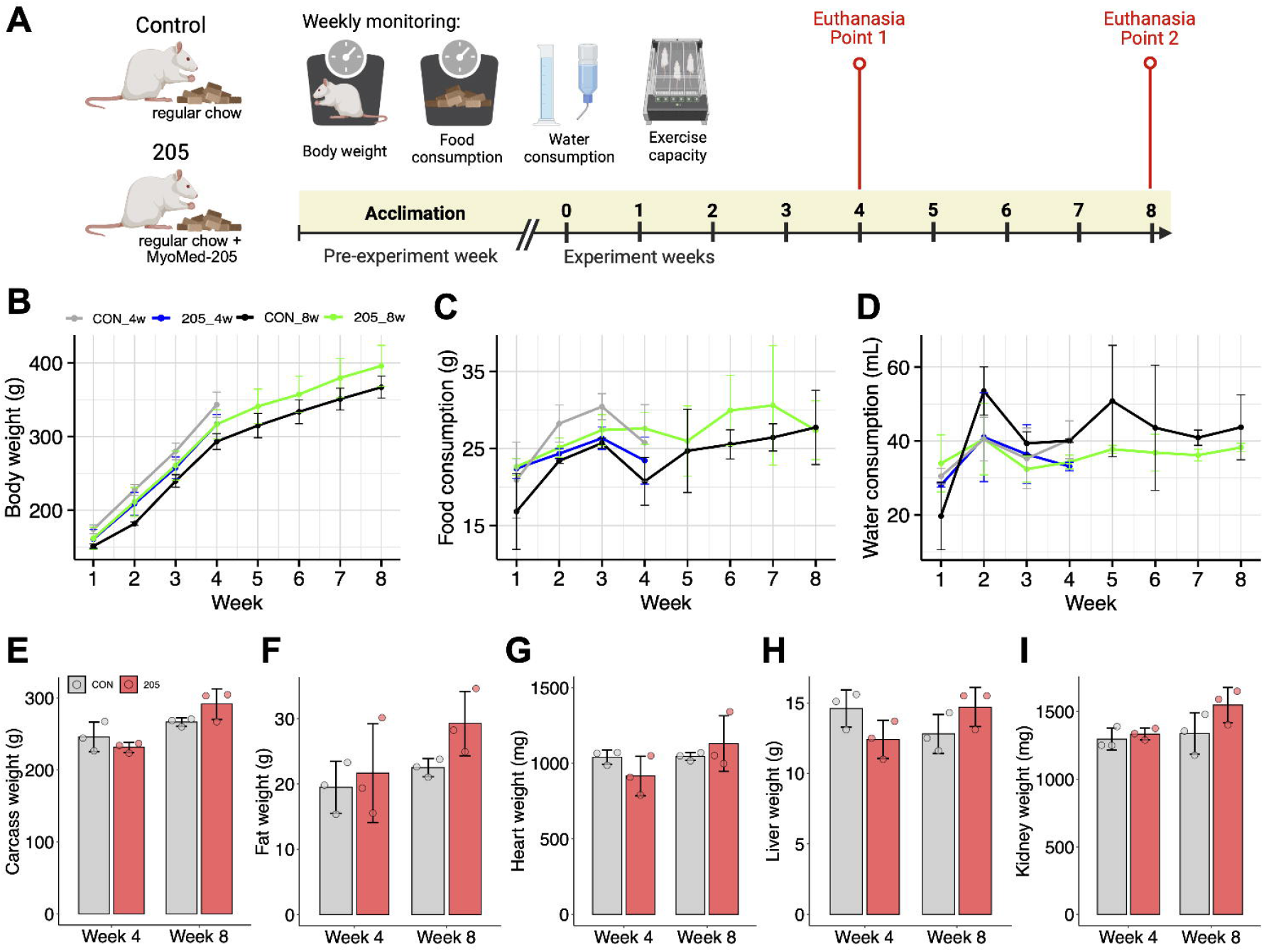
MyoMed-205 does not affect basic survival-related behaviors, body and vital organs growth after 8 weeks of administration in rats. (A) Illustrative scheme of the study experimental design (Created with BioRender.com). Time-course analysis of rats (B) Body weight, (C) Food chow intake, and (D) Water intake. Morphometric analysis of (E) Carcass weight, (F) Fat weight, (G) Heart weight, and (H) Liver weight at the 4^th^ and 8^th^-week experimental endpoints, respectively.

At each experimental endpoint, rat body composition and organ weights were determined. MyoMed-205-treated animals showed a trend to relatively higher (+10%) carcass weight (i.e., sum of skeletal muscle, bone and skin weight) after 8 weeks compared with controls (CON vs 205: 267 ± 5.7 vs 292 ± 21.2, p = 0.17, n = 3), which failed to reach statistical significance (Figure 1E). Furthermore, MyoMed-205-treated rats also showed a trend to increased (+30%) body fat mass after 8 weeks compared with controls (CON vs 205: 22.5 ± 1.38 vs 29.2 ± 4.93, p = 0.13, n = 3), although also failing to reach statistical significance (Figure 1F). Moreover, no significant differences were observed in vital organs’ weight (i.e., heart, liver and kidney) in MyoMed-205-treated rats compared with controls (Figures 1G-1I).

### MyoMed-205 enhances exercise capacity after 6 to 8 weeks of administration in rats

To explore whether MyoMed-205 long-term administration affects rat functional capacity, we performed weekly exercise capacity tests over 8 weeks using a rodent treadmill (Figure 1A). After 4 weeks, no significant changes in peak speed, running duration or distance covered were observed following MyoMed-205 administration compared with controls (CON vs 205, speed (m/min): 16.6 ± 7.6 vs 18.3 ± 5.7, p = 0.77, n = 3; duration (min): 8.7 ± 4.6 vs 9.2 ± 2.8, p = 0.88, n = 3; distance (m): 103 ± 80 vs 110 ± 60, p = 0.91, n = 3). In contrast, MyoMed-205-treated rats showed significantly enhanced exercise capacity after 6 to 8 weeks of administration compared with controls (CON vs 205, pooled comparison of mean exercise capacity between weeks 6 - 8, speed (m/min): 20.5 ± 0.9 vs 26.1 ± 1.9, p = 0.02, n = 3; duration (min): 10.3 ± 0.9 vs 13.5 ± 1.1, p = 0.01, n = 3; distance (m): 130 ± 18 vs 201 ± 28, p = 0.02, n = 3) (Figures 2A-2C).

**Figure 2.**
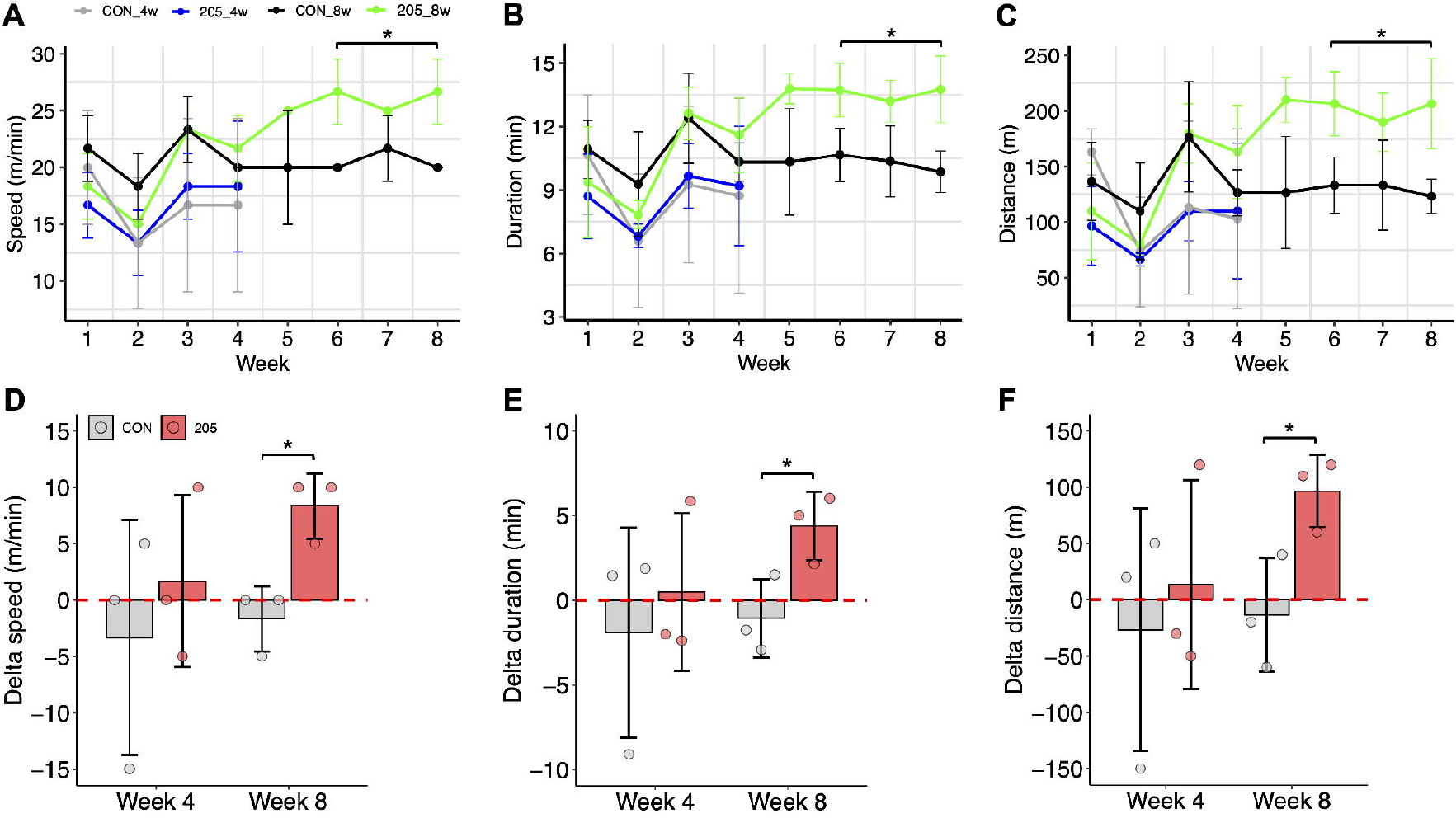
MyoMed-205 enhances exercise capacity after 6 to 8 weeks of administration in rats. Time-course analysis of rat aerobic exercise capacity indicated by the (A) Speed (m/min), (B) Duration (min), and (C) Distance (m). Average relative exercise capacity change after 4 and 8 weeks of treatment compared with intra-group baseline functional capacity for the (D) Speed (m/min), (E) Duration (min), and (F) Distance (m). *, p < 0.05 according to Welch’s t-test (n = 3/group).

At the end of the 8th week, control rats did not exhibit changes in exercise capacity compared with basal performance determined in the first week, including running speed (-8%, p = 0.42, n = 3), duration (-10%, p = 0.34, n = 3), and distance covered (-10%, p = 0.59, n =3). On the other hand, MyoMed-205-treated rats showed substantial relative gains in exercise capacity compared with the basal performance determined in the first week of administration, including increased running speed (+45%, p = 0.01, n = 3), duration (+47%, p = 0.03, n = 3), and distance covered (+87%, p = 0.04, n = 3) (Figures 2D-2F).

### MyoMed-205 does not affect soleus muscle morphometric properties after 8 weeks of administration in rats

Next, we investigated whether MyoMed-205 mediated improvements in rat exercise capacity were linked to modifications in skeletal muscle structure. For this, soleus muscle cryosections were stained with hematoxylin and eosin (H&E) for initial qualitative histological evaluation.

No significant changes were noticed in the muscle qualitative aspects (e.g., polygonal-shaped cells, peripheral nuclei number, or evidence of edema or inflammatory cells infiltrate, and regular extracellular matrix space) after 4 and 8 weeks of MyoMed-205 administration compared with controls (Figure 3A).

**Figure 3.**
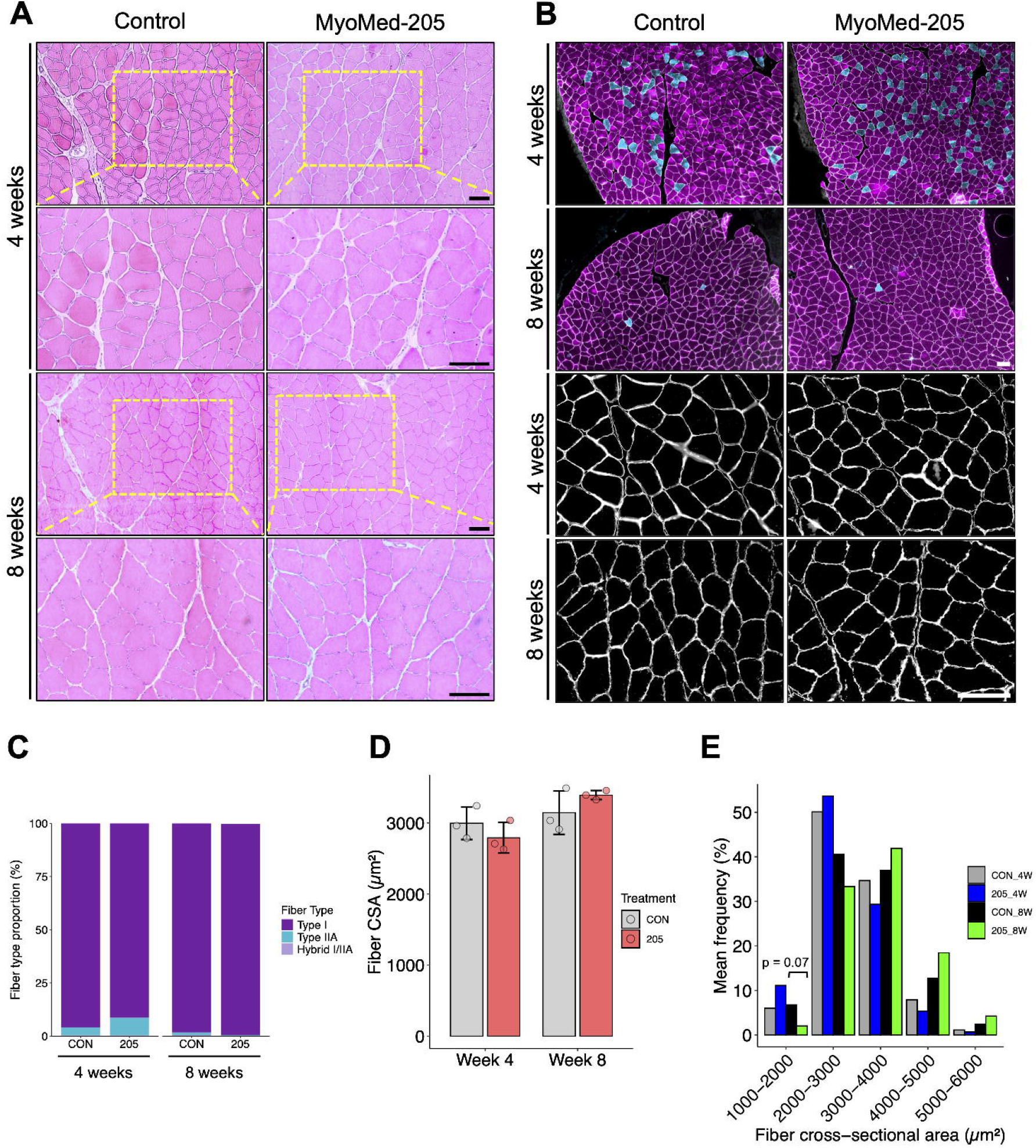
MyoMed-205 does not affect the soleus muscle histomorphometric properties after 8 weeks of administration in rats. (A) Representative photomicrographs of the soleus muscle stained with hematoxylin and eosin (H&E) for qualitative histological properties analysis after 4 and 8 weeks of administration (Scale bar = 100 μm). (B) Representative photomicrographs of the soleus muscle stained with antibodies against myosin heavy chain isoforms (Type I fiber = purple staining; Type IIa fiber = light blue staining), and dystrophin (white staining) for labelling muscle cell membrane used for fiber typing and quantitative morphometric analysis, respectively (Scale bar = 100 μm). Soleus muscle (C) fiber type relative distribution (%); (D) fiber cross-sectional area (CSA); and (E) fiber size relative distribution after 4 and 8 weeks of MyoMed-205 administration compared with controls. Statistical comparisons between groups according to Welch’s t-test (n = 3/group).

Next, we performed an immunohistochemistry assay for characterizing the soleus muscle by fiber type according to the predominant MyHC expressed, and determined their relative distribution (Figure 3B). After 4 weeks, the fiber type proportions observed were comparable between MyoMed-205-treated and control animals (CON vs 205, %, Type I: 96.0 ± 3.57 vs 91.4 ± 7.29; Type IIa: 3.74 ± 3.48 vs 8.28 ± 7.0; and hybrid I/IIa: 0.30 ± 0.53 vs 0.31 ± 0.37). Similarly, the fiber type distribution was not altered after 8 weeks in MyoMed-205-treated compared with controls (CON vs 205, %, Type I: 98.3 ± 1.83 vs 99.1 ± 0.77; Type IIa: 1.27 ± 1.37 vs 0.50 ± 0.70; and hybrid I/IIa: 0.45 ± 0.46 vs 0.10 ± 0.12). These data confirmed that the rat soleus muscle is predominantly composed of slow-twitch type I fibers (>90%) at both experimental endpoints evaluated in this study (Figure 3C).

Given the very low percentage of fast-twitch fibers in the rat soleus muscle, we performed a quantitative morphometric analysis of the muscle fibers using cryosections labeled with the membrane-located protein, dystrophin. In this context, no significant changes were found in the fibers’ CSA in MyoMed-205-treated animals compared with controls after 4 weeks of administration (CON vs 205, 2997 ± 230 vs 2791 ± 215 μm^2^, p = 0.32, n = 3). Similarly, MyoMed-205-treated rats showed no significant increase in soleus muscle fibers CSA compared with controls (CON vs 205: 3145 ± 305 vs 3392 ± 64, p = 0.29, n = 3) (Figure 3D).

Fiber size distribution analysis was carried out to investigate MyoMed-205’s effect on soleus muscle fiber morphometric properties over time. After 4 weeks, no changes in the fiber distribution pattern were observed in MyoMed-205-treated rats compared with controls. After 8 weeks, MyoMed-205-treated animals showed a trend for lower percentage of small fibers compared with controls (e.g., CON vs 205, 1000 – 2000 μm^2^: 7% vs 2%, p = 0.07, n = 3). Concomitantly, a relative, but not statistically significant, higher cumulative percentage of muscle fibers with bigger CSA was found in MyoMed-205-treated rats compared with controls (e.g., CON vs 205, 3000 – 5000 μm^2^, 50% vs 60%, p = 0.32, n =3) (Figure 3E).

### MyoMed-205 does not affect soleus intramuscular lipid content after 8 weeks in rats

To examine whether MyoMed-205 long-term administration might affect intramuscular lipid accumulation, we performed the Oil Red O (ORO) staining of the soleus muscle cryosections from MyoMed-205-treated and control rats (Figure 4A). After 4 weeks, we found that MyoMed-205-treated rats showed lower intramuscular lipid content compared with controls (CON vs 205, average ORO area (%): 4.0 ± 0.8 vs 1.9 ± 0.3, p = 0.03, n = 3) (Figure 4B). However, this effect was not further observed after 8 weeks, when MyoMed-205-treated rats showed comparable levels of soleus intramuscular lipid content with controls (CON vs 205, average ORO area (%): 1.8 ± 0.2 vs 1.6 ± 0.4, p = 0.66, n = 3) (Figure 4C).

**Figure 4.**
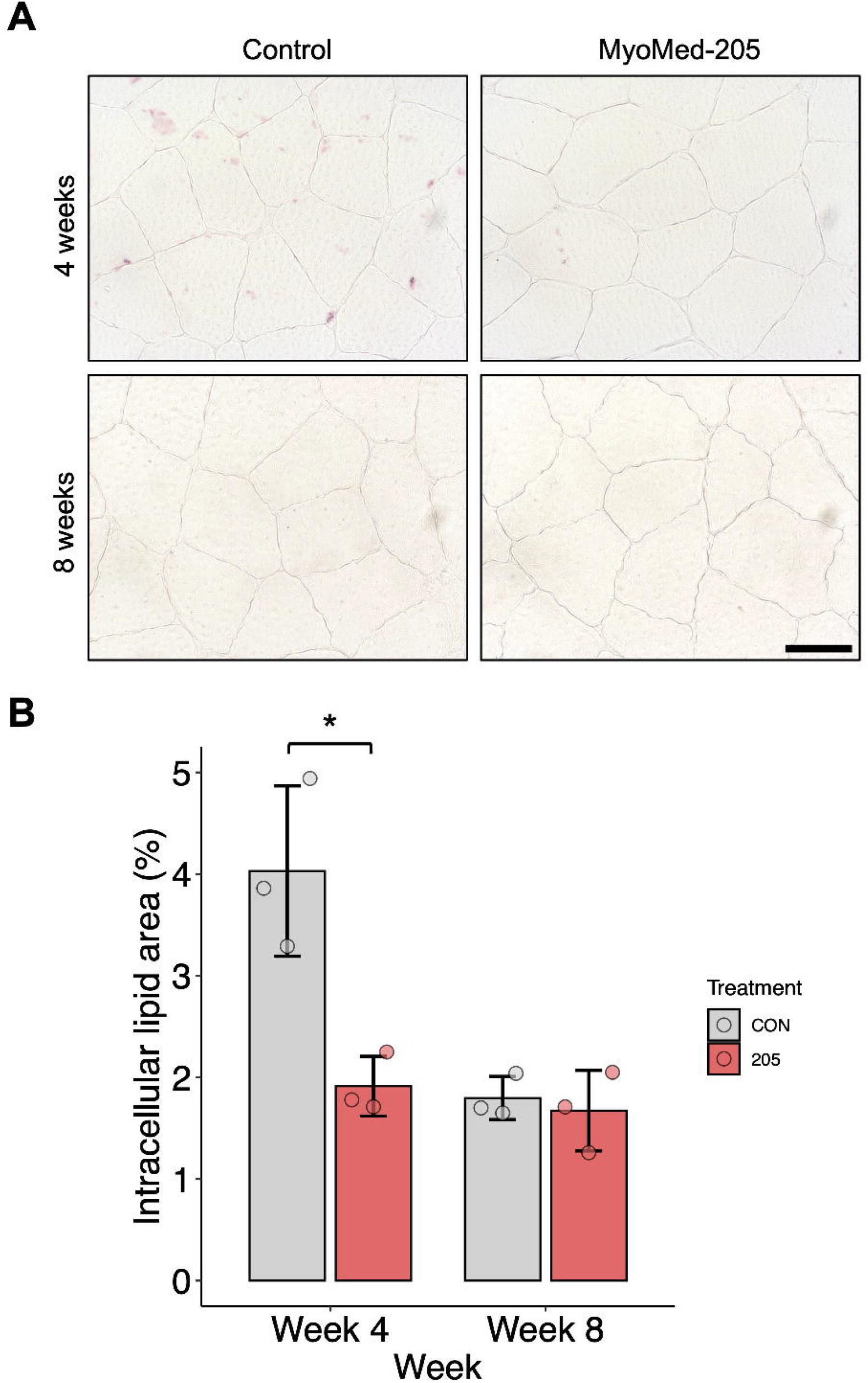
MyoMed-205 does not affect the soleus muscle intramuscular lipid deposit after 8 weeks of administration in rats. Oil Red O staining was used to assess the intramuscular lipid levels in the soleus muscle. (A) Representative Oil Red O staining photomicrographs of soleus muscle (Scale bar = 50 μm). (B) Intracellular lipid area after 4 weeks and 8 weeks of administration. *, p < 0.05 according to Welch’s t-test (n = 3/group).

### MyoMed-205 does not affect soleus intramuscular collagen content after 8 weeks in rats

To assess whether MyoMed-205 long-term administration might affect intramuscular collagen deposition and extracellular matrix (ECM) structure, we performed the Picrosirius Red (PSR) staining of soleus muscle samples collected from MyoMed-205-treated and control rats (Figure 5A). After 4 weeks, no differences in collagen muscle content between MyoMed-205-treated and control animals (CON vs 205, collagen (%): 15.0 ± 4.5 vs 11.2 ± 1.4, p = 0.28, n = 3). Likewise, MyoMed-205-treated rats showed similar soleus muscle collagen content compared with controls after 8 weeks of administration (CON vs 205, collagen (%): 11.0 ± 3.0 vs 11.9 ± 2.4, p = 0.68, n = 3) (Figure 5B).

**Figure 5.**
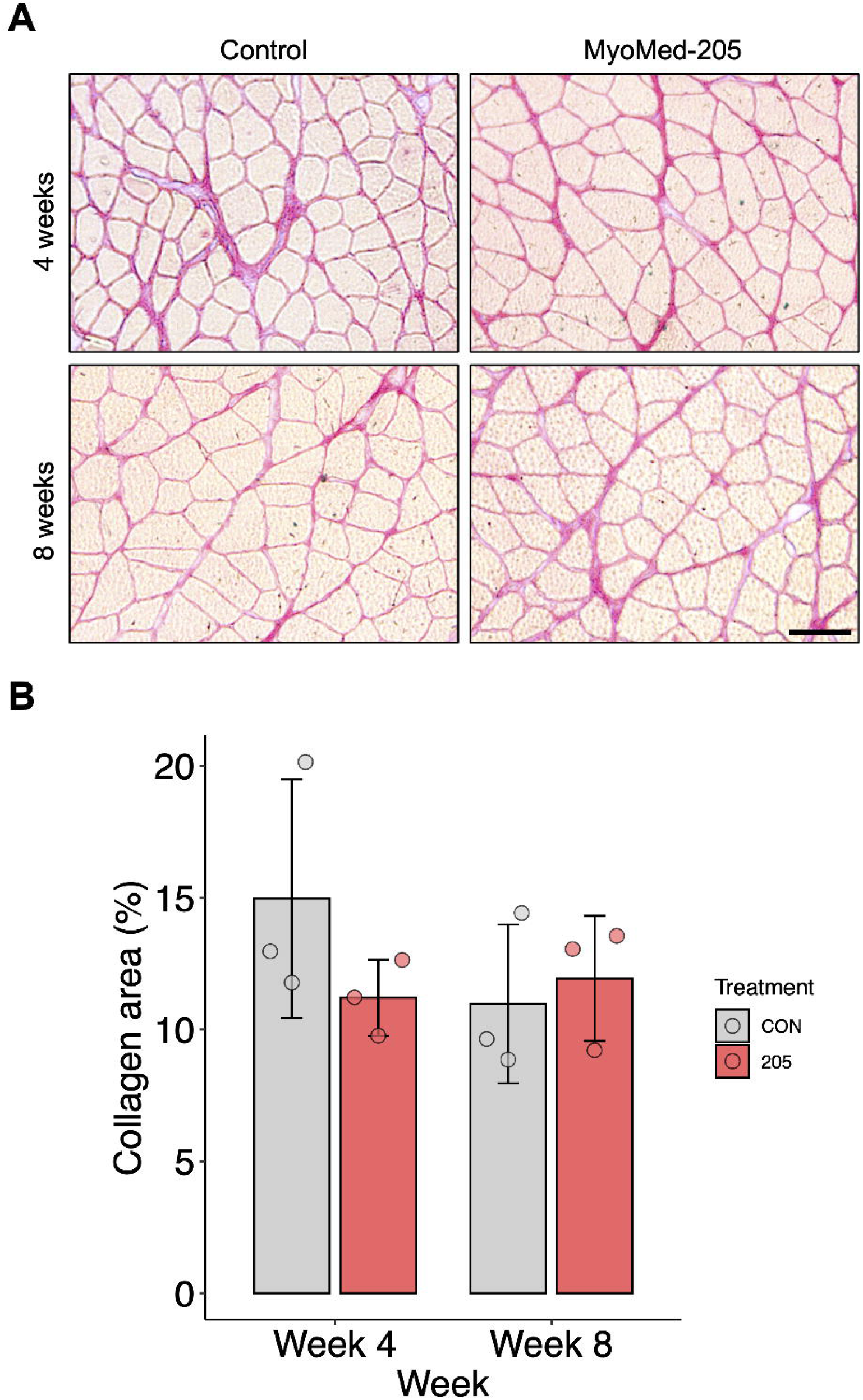
MyoMed-205 does not affect soleus muscle collagen content after 8 weeks of administration in rats. Picrosirius Red staining was used to assess the collagen deposit in the soleus muscle. (A) Representative Picrosirius Red staining photomicrographs of soleus muscle (Scale bar = 100 μm). (B) Collagen relative content in the soleus muscle after 4 and 8 weeks of administration (n = 3/group).

## Discussion

Maintenance of skeletal health is essential for long-term functional independence, quality of life and healthspan. At the molecular level, this process is linked to the muscle-specific E3 ubiquitin ligase MuRF1, a known key regulator of muscle mass, function and metabolism. The development of pharmacological MuRF1 inhibitors, such as the small molecule MyoMed-205, has advanced the investigation of the translational potential of MuRF1-targeting compounds to counteract muscle wasting and weakness under catabolic or disease conditions. However, despite these recent advances, the long-term effects of MuRF1 inhibition on growth, behavior, and muscle physiology under basal conditions remain unknown. In this exploratory study, we assessed morphological, functional, and skeletal muscle structural properties in animals subjected to long-term treatment with MyoMed-205. Our findings indicate that long-term MyoMed-205 administration supports normal somatic growth while enhancing functional exercise capacity independently of significant remodeling in the soleus muscle structure in rats.

We initially investigated whether long-term MyoMed-205 administration affects normal development in rats under basal conditions. In this context, MyoMed-205 did not alter body weight development, basic survival-related behaviors (e.g., food and water intake), or vital organ weight, including heart, liver or kidneys. These findings are consistent with previous studies showing that MuRF1 knockout mice are viable, fertile, and exhibit normal muscle development and lifespan compared with wild-type animals ^13,14^. Previous acute toxicological studies from our group support that even high doses of MyoMed-205 (300–2000 mg/kg body weight) do not provoke behavioral abnormalities or gross organ macroscopic changes in rats ^22^. Collectively, these data indicate that long-term MyoMed-205 administration does not impair somatic or vital organ development in rats under basal conditions.

MuRF1 is known to play a key role in the regulation of myofibrillar protein function and degradation ^6,13,26^. Thus, we monitored rat exercise capacity over time and interestingly found that MyoMed-205-treated rats exhibited enhanced exercise capacity, including substantial gains in the peak speed, running endurance, and distance covered after 6 to 8 weeks of administration. Notably, the magnitude of the functional gains observed herein is comparable to those reported in experimental studies following 4 to 8 weeks of moderate-to-high intensity aerobic exercise training in rats ^27–29^. While it is known that aerobic exercise training under pathological conditions confers protective health effects partly through downregulation of MuRF1 in humans ^30,31^, the direct impact of MuRF1 genetic or pharmacological blockade on functional capacity has not been previously demonstrated. In this line, previous findings suggest that MuRF1 deletion increases absolute muscle force in mice under basal conditions ^11^. Although increased muscle strength does not necessarily translate into improved aerobic exercise capacity, it is reasonable to hypothesize that MyoMed-205-mediated increases in functional capacity reported here could involve improved skeletal muscle contractile function.

Despite enhancing functional capacity, long-term MyoMed-205 administration seems not to affect overall musculoskeletal mass (e.g., carcass weight) or soleus muscle fiber CSA. These findings are consistent with previous reports suggesting only small or undetectable effects of MuRF1 deletion on skeletal muscle mass in mice under basal conditions ^13,32^. Taken together, these evidences suggest that MuRF1 does not represent a sufficient stimulus alone to regulate skeletal muscle mass under physiological circumstances, although it reportedly plays a crucial role in muscle mass control under catabolic stress ^9,10,14,33^. Nevertheless, as suggested herein, targeting MuRF1 with small molecules can significantly improve exercise capacity. Based on previous mechanistic findings ^13,14,23,34^, we conjecture that the functional gains described here could likely be linked to MuRF1-mediated regulation of muscle function or energy metabolism pathways rather than from direct hypertrophic adaptations.

In addition to its role in myofibrillar protein turnover, MuRF1 also regulates muscle energy metabolism ^13,14,34,35^. In the present study, we observed a trend toward increased body fat mass in rats after 8 weeks of MyoMed-205 administration. Consistent with this finding, MuRF1 knockout mice display altered lipid metabolism, characterized by elevated serum triglycerides, increased basal energy expenditure, and preferential usage of lipid oxidation for cellular energy metabolism ^21,36^. Furthermore, MuRF1 deficiency has been linked to increased hepatic lipid deposition in mice under dexamethasone-induced stress ^37^. In the present study, soleus intramuscular lipid content was found to be decreased or unaltered after 4 and 8 weeks of MyoMed-205 administration, respectively. These observations are consistent with our recent study ^23^ demonstrating that MyoMed-205 could downregulate pro-adipogenic mechanisms and restrict intramuscular lipid accumulation, such as in the diaphragm under acute denervation. Altogether, these findings suggest that disruption of MuRF1 expression/activity may affect systemic lipid metabolism, which could contribute to augmented body fat mass linked to decreased intramuscular lipid deposition under basal conditions.

MuRF1 has also been implicated in the regulation of extracellular matrix (ECM) remodeling during muscle atrophy. In this context, genetic or pharmacological inhibition of MuRF1 downregulates the transcription of key ECM components (e.g., collagen) under stress induced by dexamethasone and denervation ^23,38^. Furthermore, in experimental models of heart failure, MyoMed-205-mediated downregulation of muscle fibrosis has been associated with improved functional outcomes ^17^. In contrast, our findings herein suggest that long-term MyoMed-205 administration did not affect soleus muscle collagen content under basal conditions. Thus, we conjecture that MuRF1-dependent regulation of muscle ECM could primarily be involved in the control of muscle ECM remodeling under catabolic conditions rather than under basal conditions.

These findings should be interpreted in light of the following study limitations: First, the small sample size (n = 3/group) reflects the exploratory nature of this long-term study and adherence to ethical principles of animal use reduction for experimentation. Nevertheless, to our knowledge, this is the first investigation to systematically explore the long-term effects of small-molecule targeting MuRF1 on development, behavior, systemic functional capacity and muscle physiology under basal conditions. Second, the exclusive use of male rats in this study limits the direct extrapolation of these findings to females and insights into sex-dependent effects. Third, the assessment of a single saturating-level dosage of MyoMed-205 restricts potential inferences on dose-dependent effects and warrants forthcoming studies. Fourth, direct measurements of muscle function were not performed, precluding definitive conclusions regarding the long-term effect of MyoMed-205 on muscle mechanical properties. Finally, histological and morphometric analyses were restricted to a single muscle (i.e., soleus, slow-twitch properties), and effects in other muscle types with distinct functional or metabolic properties remain to be determined.

## Conclusion

Long-term treatment with a small-molecule targeting MuRF1 (i.e., MyoMed-205) enhances functional exercise capacity in rats under basal conditions. Furthermore, during the 8-week observation period, MyoMed-205 has not adversely affected body growth, basic survival-related behaviors such as food and water intake, or soleus muscle structural properties (e.g., fiber type distribution, size, as well as intramuscular lipid content and collagen content). These findings suggest that small molecules targeting MuRF1 could improve functional capacity without disrupting normal growth or muscle structure under physiological circumstances. Collectively, our findings indicate the potential of MuRF1-targeting pharmacological strategies to improve functional capacity without impairing skeletal muscle structure. Future studies are warranted to determine whether these effects extend to skeletal muscles with distinct functional and metabolic profiles, to characterize sex-specific responses, dose dependency effects, and the underlying molecular mechanisms.

## Author’s Contributions (CRediT)

### Fernando Ribeiro

Conceptualization, Methodology, Investigation, Formal analysis, Data Curation, Visualization, Writing – Original Draft, and Writing – Reviewing and Editing; **Luis D. Chinait:** Investigation and Writing – Reviewing and Editing. **Maria Rita C. Rodrigues:** Investigation and Writing – Reviewing and Editing; **Siegfried Labeit:** Funding Acquisition, Resources, Supervision, and Writing – Reviewing and Editing; **Anselmo S. Moriscot:** Conceptualization, Methodology, Funding Acquisition, Resources, Project Management, Supervision, Writing – Original Draft, and Writing – Reviewing and Editing.

## Acknowledgements

We thank Kelly P. Borges for her excellent laboratory technical assistance.

## Data Availability Statement

The data supporting the findings of this study are included in the article. Additional information is available from the corresponding author upon reasonable request.

